# Androgen Receptor is a Determinant of Melanoma targeted drug resistance

**DOI:** 10.1101/2022.05.27.493720

**Authors:** Anastasia Samarkina, Markus Kirolos Youssef, Paola Ostano, Soumitra Ghosh, Min Ma, Beatrice Tassone, Tatiana Proust, Giovanna Chiorino, Mitchell P. Levesque, Gian Paolo Dotto

## Abstract

Melanoma provides a primary benchmark for targeted drug therapy. Most melanomas with BRAF^V600^ mutations regress in response to BRAF/MEK inhibitors (BRAFi/MEKi). However, nearly all relapse within the first two years, and there is a connection between pathways involved in BRAFi/MEKi-resistance and poor response to immune checkpoint therapy. We recently showed that androgen receptor (AR) activity is required for melanoma cell proliferation and tumorigenesis. Here we find that AR expression is markedly increased in BRAFi resistant melanoma cells as well as in sensitive cells soon after BRAFi exposure. Increased AR expression is by itself sufficient to render melanoma cells BRAFi-resistant, eliciting transcriptional changes of BRAFi resistant subpopulations and elevated *EGFR* and *SERPINE1* expression of likely clinical significance. Inhibition of *AR* expression and activity blunts changes in gene expression and suppresses proliferation and tumorigenesis of BRAFi-resistant melanoma cells, enhances MHC I expression and CD8+ T cells infiltration. Our findings point to targeting AR as a possible co-adjuvant approach for the prevention and management of the disease.

## Introduction

Significant differences exist in melanoma mortality between men and women across all ages and after adjusting for tumor variables (Breslow thickness, histologic subtypes, body site, and metastatic status) ^1^. As for sexual dimorphism in other cancer types ^2^, even for melanoma, differences in sex hormone levels and/or downstream pathways are likely to play a role ^3^. Sex hormone signaling can affect cancer susceptibility through multiple intrinsic and extrinsic mechanisms, impacting cancer stem cell renewal, the tumor microenvironment, the immune system, and the overall metabolic balance of the organism ^2,4-6^. As early as 1980, it was proposed that differences in androgen levels could help explain the lower survival of male versus female melanoma patients ^7^. Recent epidemiological evidence links elevated free testosterone levels in male human populations with a high risk of melanoma as the only other cancer type besides prostate ^8^.

In our recent work, we have found that the androgen receptor (AR) is heterogeneously expressed in melanoma cells, both at the single-cell intralesional level and among lesions at various stages of the disease ^9^. Irrespective of expression levels, silencing of the *AR* gene and pharmacological inhibition of AR activity suppresses proliferation and induces cellular senescence of a relatively large panel of melanoma cells from both male and female patients ^9^. AR plays an essential function in this context by bridging the transcription and DNA repair machinery, maintaining genome integrity. In both cultured melanoma cells and tumors *in vivo, AR* gene silencing or treatment with AR inhibitors leads to chromosomal DNA breakage in the absence of other exogenous triggers, leakage into the cytoplasm, STING activation, and a STING-dependent pro-inflammatory cascade ^9^.

In the present study, we have assessed the translational significance of suppressing AR signaling in the context of melanoma response to targeted drug treatments, specifically BRAF inhibitors. ∼50% of all melanomas harbor BRAF^V600^ mutations, with >90% of these expressing the V600E or K amino acid substitutions. Although >80% of patients with BRAF^V600E/K^ melanomas initially respond to highly specific BRAF and MEK inhibitors (BRAFi/MEKi), nearly all patients relapse after seven months to two years ^10^. Most BRAFi/MEKi-resistant melanomas are also resistant to immunotherapies ^11^, with a cancer cell-instructed mechanism that does not depend on selection by the immune system ^12,13^. Initial treatment of melanoma patients with BRAFi/MEKi elicits recruitment and activation of immune cells ^14^, similarly to what we found in mouse xenografts with melanoma cells with *AR* gene silencing or inhibition ^9^. In melanomas with acquired BRAFi/MEKi resistance, an opposite modulation of the immune cell response occurs, which can be attributed, in part, to epigenetic/transcriptional regulatory changes that have the potential of being pharmacologically reversed ^14^.

We show that increased AR expression and activity are part of the response of melanoma cells with BRAF^v600^ mutations to treatment with BRAF inhibitors and that increased AR expression is sufficient to render these cells resistant to these drugs, inducing transcriptional changes of BRAFi resistant subpopulations of likely clinical significance. Conversely, treatment with AR inhibitors suppresses proliferation and tumorigenicity of BRAFi resistant melanoma cells, enhancing CD8+ T cells infiltration. Our findings raise the exciting possibility that targeting AR signaling, which is a routine treatment of metastatic prostate cancer, can also enhance the efficacy of melanoma targeted therapy.

## Results

### 1. BRAFi treatment induces AR expression in melanoma cells

Acquisition of BRAFi resistance by melanoma cells can be a dynamic process that is induced in culture by the drug treatment ^15,16^. Treatment of primary human melanoma cells (M121224) with multistep increases of the BRAF inhibitor Dabrafenib (DAB) resulted in the emergence of cells actively proliferating in the presence of this compound. RT-qPCR and immunoblot analysis showed substantially increased AR expression already at lower doses (Fig. 1A). A consistent increase in AR expression was found in additional primary and established melanoma cells selected for BRAFi resistance by immunoblot and immunofluorescence analysis as well as RT-qPCR (Fig. 1B, C, Suppl. Fig. 1A). While AR expression was upregulated in all BRAFi-resistant cell lines relative to parental cells, other genes connected with the acquisition of BRAFi resistance, such as *MITF, SOX9, SOX10, ZEB1*, and *ZEB2* ^17^ were more unevenly modulated (Fig. 1D; Suppl. Fig. 1B, C). Upregulation of AR expression was also found in the clinical setting, by immunofluorescence analysis of matched lesions arising in the same patients before and after BRAFi/MEKi therapy (Suppl. Fig. 2).

**FIGURE 1:**
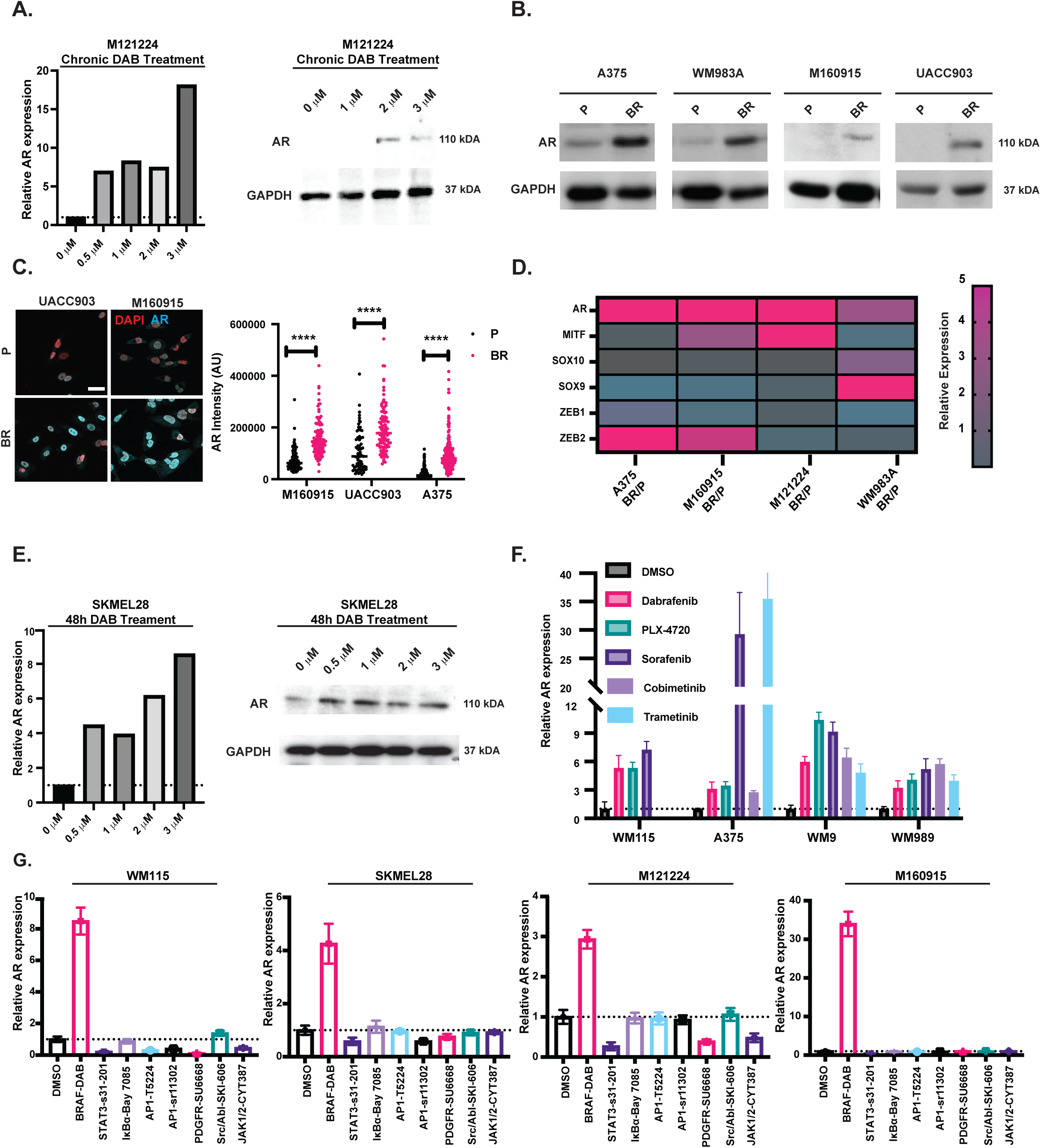
BRAFi treatment of melanoma cells results in increased AR expression. A) RT-qPCR and immunoblot analysis of AR expression in primary human melanoma cells (M121224) cultured with multistep weekly increases of the BRAF inhibitor Dabrafenib (0.5, 1, 2, and 3 µM). Cells were collected at the end of each week of treatment and analyzed together with the untreated parental cells for levels of AR expression by RT-qPCR, with *RPLP0* for internal normalization, and immunoblotting, with GAPDH as an equal loading control. Results of similar independent experiments with additional cell lines are shown in Suppl. Fig. 1A. B) Immunoblot analysis of AR expression in additional primary (M160915) and established melanoma cell lines (A375, WM983A, and UACC903) selected for BRAFi resistance (BR) by multistep cultivation in increasing amounts of Dabrafenib (up to 3 µM) as in the previous panel, versus untreated parental cells (P). Immunoblotting for GAPDH was used as an equal loading control. C) Immunofluorescence analysis of AR expression in primary and established melanoma cells selected for BRAFi resistance (BR) as in the previous panels versus untreated parental cells (P). Shown are representative images and quantification of AR nuclear intensity signal in arbitrary units (AU) per cell (dots) together with a mean, examining >100 cells per sample, unpaired *t*-test, **** p<0.0001. Color scale: red, DAPI; cyan, AR. Scale bar: 40μm. D) Relative expression of the indicated genes in BRAFi resistant (BR) primary (M160915, M121224) and established melanoma cells (A375 and WM983A) versus parentals (P). Results are represented as a heatmap of changes in gene expression as assessed by RT-qPCR analysis with *RPLP0* for internal normalization. Individual bar plots of the results are shown in Supplementary Fig. 1B. Magenta and grey: up- and down-regulated genes, respectively. E) AR expression in melanoma cells (SKMEL28) for 48 hours with Dabrafenib (0.5, 1, 2, and 3 µM) versus DMSO control. Cells were analyzed by RT-qPCR, with *RPLP0* for internal normalization, and immunoblotting, with GAPDH as an equal loading control. F) AR expression in the indicated melanoma cells at 48 hours of treatment with various BRAF (Dabrafenib, PLX-4720, and Sorafenib; 0.5 µM) and MEK (Cobimetinib and Trametinib; 5 nM) inhibitors versus DMSO control. RT-qPCR results are expressed as fold changes relative to untreated controls, after *RPLP0* normalization. Results of similar independent experiments with these and additional cell lines are shown in Suppl. Fig. 1D. G) AR expression in the indicated primary and established melanoma cells at 48 hours of treatment with inhibitors of the indicated molecules/pathways along with corresponding chemical names at concentrations specified in methods. RT-qPCR results are expressed as fold changes relative to untreated controls, after *RPLP0* normalization.

**FIGURE 2:**
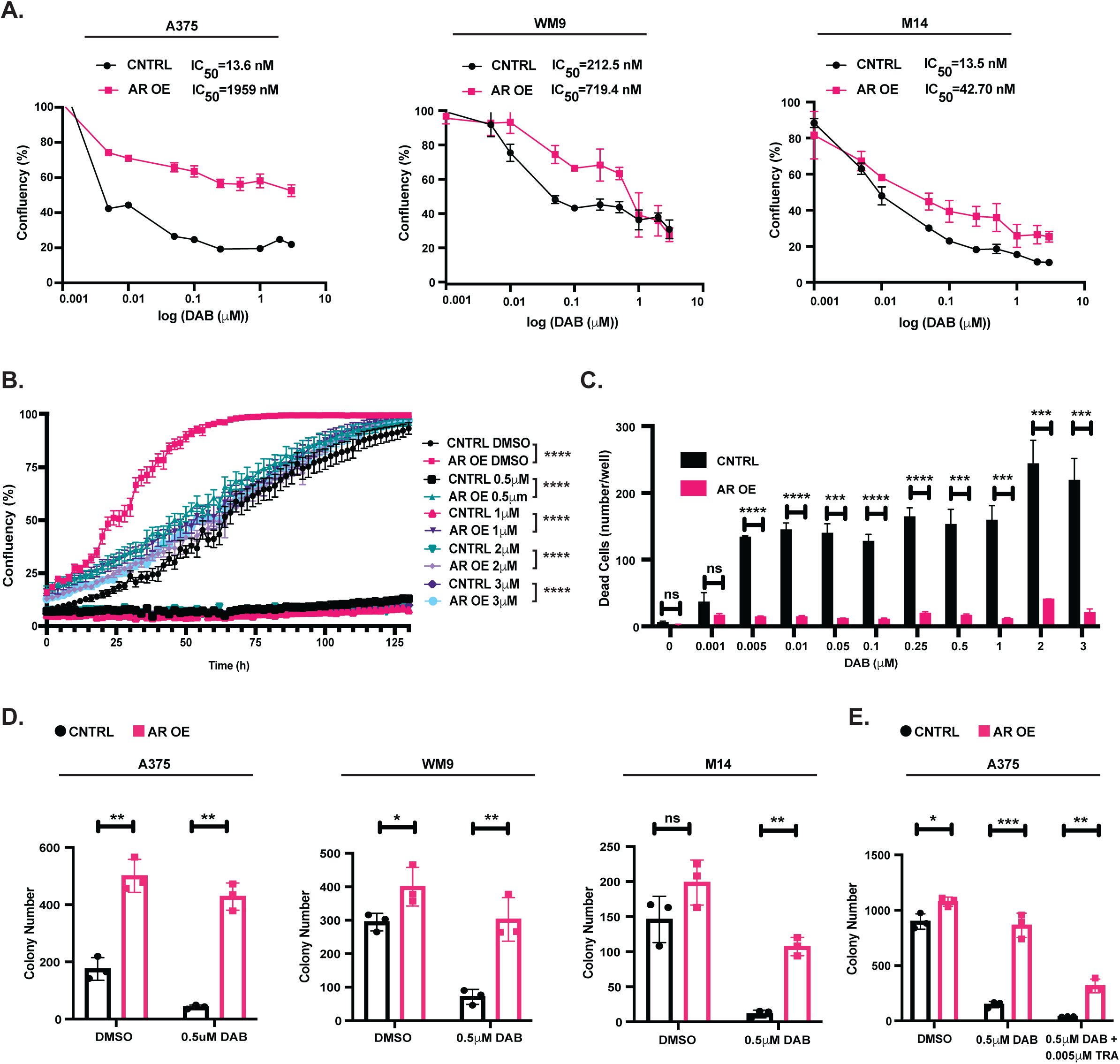
AR overexpression confers BRAFi resistance. A) Cell density assays (CellTiter-Glo) of the indicated melanoma cells stably infected with an AR overexpressing lentivirus (AR OE) versus LacZ expressing control (CNTRL) and treated with the indicated increasing amounts of Dabrafenib for 72 hours. For each condition, cells were tested in triplicate dishes, and results are expressed relative to DMSO control. The calculated IC_50_ for each condition is indicated above. B) Proliferation by live-cell imaging assays (IncuCyte) of AR overexpressing (AR OE) versus control (CNTRL) A375 melanoma cells (obtained as in the previous panel) cultured with the indicated concentrations of Dabrafenib versus DMSO. Cells were plated in triplicate wells in 96-well plates followed by cell density measurements (four images per well every 4 h for 128 h). cultures, n = 3; Pearson r correlation test. ****, p< 0.0001. C) BRAFi-induced cell death as detected by live-cell staining (IncuCyte, Cytotox Red) of AR overexpressing (AR OE) versus control (CNTRL) melanoma cells (A375) at 72 hours of treatment with Dabrafenib at the indicated concentrations as in the previous panel. Four images per well cultures, n (cultures) = 3; unpaired *t*-test, ns, non-significant, ***, p< 0.001; ****, p<0.0001. D) Clonogenicity assays of the indicated melanoma cells transduced with an AR overexpressing lentivirus (AR OE) versus empty vector control (CNTRL) treated with Dabrafenib (DAB, 0.5µM) versus DMSO. Cells were plated in triplicates at clonal density (5000 cells / 6 cm dish) followed by 1-week cultivation. Macroscopically detectable colonies were counted after crystal violet staining. n(dishes)=3, unpaired *t*-test, *, p<0.05; **, p<0.01, ns, non-significant. E) Clonogenicity assays of AR-overexpressing versus control A375 melanoma cells as in the previous panel treated with Dabrafenib (0.5µM) individually and in combination with the MEK inhibitor Trametinib (5nM) as in the previous panel. n(dishes)=3, unpaired *t*-test, *, p<0.05; **, p<0.01, ***, p<0.001.

AR upregulation may result from chronic BRAFi treatment or be part of an acute response. In fact, pronounced induction of *AR* expression occurred in SKMEL28 melanoma cells as well as other primary and established melanoma cells already by 48 hours of Dabrafenib treatment (Fig. 1E, Suppl. Fig. 1D). *AR* expression was specifically induced in melanoma cells by treatment with Dabrafenib and other BRAF and MEK inhibitors and not inhibitors of other key signaling pathways, such as NF-κB, STAT3, and AP-1 (Fig. 1F, G).

Expression of the *AR* gene can be positively regulated by the CREB1, c-Myc, LEF/ß-catenin, Foxo3a, Sp1, Twist, and SREBP-1 transcription factors ^18^. To gain insights into the mechanisms responsible for the induction of AR expression by Dabrafenib, we probed into the transcriptomic profiles of 3 different melanoma cell lines plus/minus treatment with this compound for 48 hours, as considered in greater detail further below. Of the positive regulators of *AR* gene transcription, *FOXO3* and *CREB1* were consistently upregulated in all three melanoma lines by Dabrafenib treatment and *SP1* in two of the three, while others were more unevenly or not modulated (Suppl. Fig. 3A, Suppl. Table 1). The connection of *FOXO3, CREB1* and *SP1* with *AR* expression was further validated by analysis of the transcriptomic profiles of a large melanoma cohort (TCGA), showing a significant correlation between expression levels of these genes and *AR* (Suppl. Fig. 3B).

**FIGURE 3:**
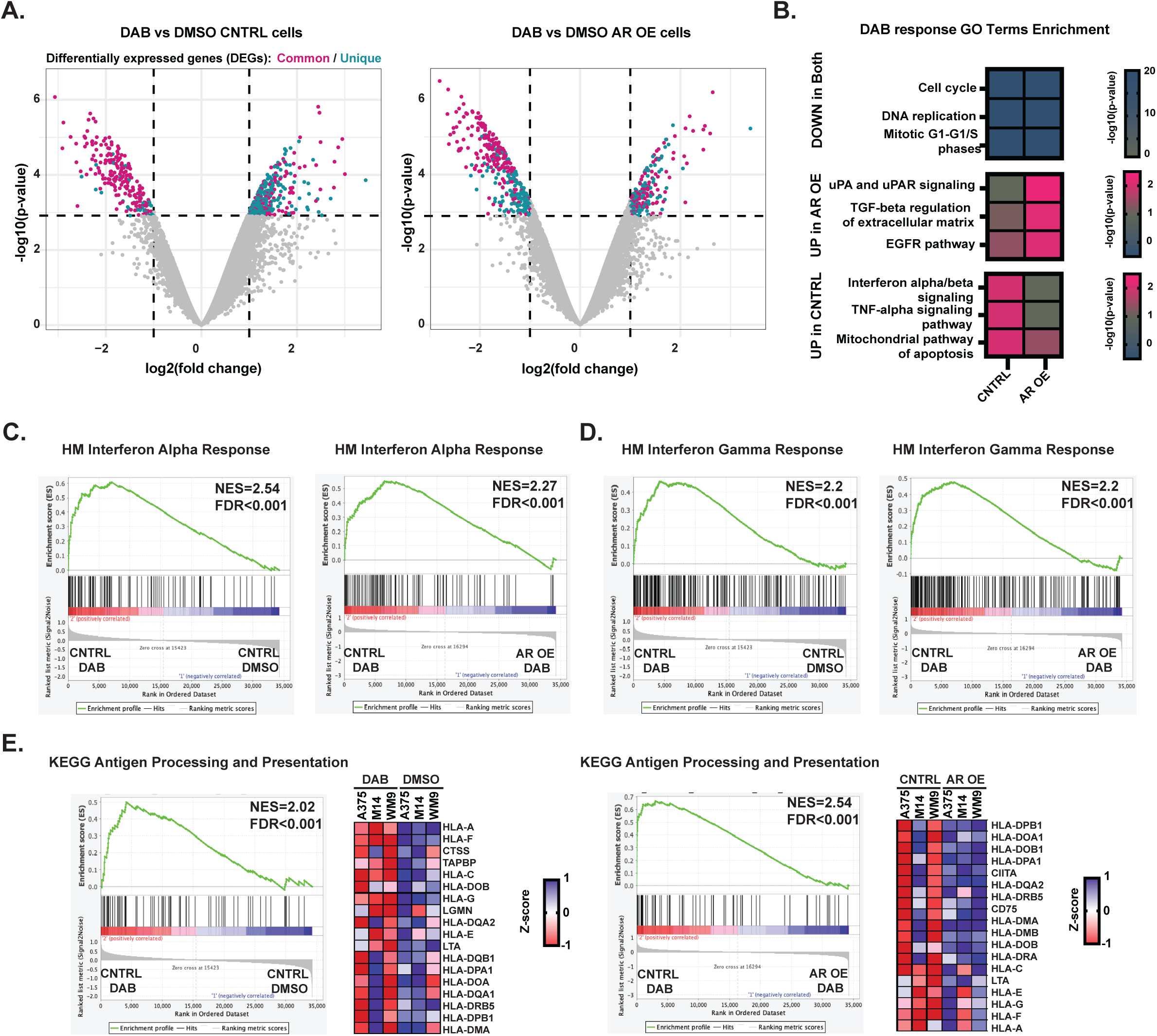
Increased AR expression perturbs the transcriptional response of melanoma cells to BRAFi. A) Transcriptional response of melanoma cells plus/minus AR overexpression to acute BRAFi treatment. Volcano plot of transcriptional changes consistently elicited in A375, M14, and WM9 melanoma cells infected with control (LacZ expressing) (left) or AR overexpressing (AR OE) (right) lentiviruses by 48 hours of treatment with Dabrafenib (0.5 µM) versus DMSO. The x-axis shows the log_2_(fold change), and the y-axis shows the −log_10_(p-value). Each dot represents one gene, with colored dots corresponding to genes with a false discovery rate threshold of < 0.05 and log fold-change threshold of -1 and 1. Magenta and cyan dots correspond to genes similarly and specifically modulated by Dabrafenib treatment in control versus AR overexpressing melanoma cells, respectively. A complete list of modulated genes in the three melanoma cell lines is provided in Suppl. Table 2. B) Functionally relevant gene ontology families significantly downmodulated by BRAFi treatment in both control and AR overexpressing cells (upper), and gene families modulated only in control (middle) or in AR overexpressing cells (bottom). The -log_10_(p-value) is indicated by the heatmap color scale. A full list of modulated gene families is provided in Suppl. Table 2. C-E) Gene Set Enrichment Analysis (GSEA) of transcriptional profiles of control melanoma cells (A375, M14, and WM9) plus/minus Dabrafenib treatment (left panel) and of Dabrafenib-treated control versus AR overexpressing cells (right panel) using predefined gene signatures of interferon alpha (C) and gamma response(D) and antigen processing and presentation (E) derived from the hallmark gene set (HM) ^42^ and KEGG ^43^ collections. Genes are ranked by signal-to-noise ratio in Dabrafenib versus DMSO treated melanoma cells; the position of individual genes is indicated by black vertical bars; the enrichment pattern is in green. In (E), GSEA and the leading-edge analysis of the antigen processing and presentation signature are shown in each of the three melanoma lines plus/minus Dabrafenib treatment (left) and of Dabrafenib-treated control versus AR overexpressing cells (right).

Thus, *AR* gene expression is consistently induced in BRAFi-resistant melanoma cells as well as in naïve melanoma cells upon acute exposure to BRAF/MEK inhibitors, with co-regulation by CREB1, Foxo3a, and Sp1 as likely involved.

### 2. Increased AR expression triggers a BRAFi resistant phenotype

To assess the functional significance of the findings, we infected three different melanoma lines (A375, WM9, and M14) with an AR overexpressing lentivirus (AR OE) versus LacZ-expressing control (CNTRL). In dose-response cell growth assays, the half-maximal inhibitory concentration (IC_50_) of Dabrafenib at 72 hours of treatment was drastically increased by AR overexpression in all three cell lines (Fig. 2A). In one-week cell imaging assays (Incucyte), the proliferation of control A375 cells was suppressed by Dabrafenib treatment at all tested concentrations. In contrast, that of AR overexpressing cells was initially reduced, but cultures eventually attained the same density as untreated controls (Fig. 2B). In parallel, Dabrafenib treatment induced cell death to a much greater extent in control than AR overexpressing cells (Fig. 2C).

The findings were expanded by clonogenicity assays. The number of colonies produced by control cells was drastically reduced by Dabrafenib treatment. AR overexpression enhanced the colony-forming ability of cells already under basal conditions and effectively counteracted the decrease caused by this compound (Fig. 2D). Similar protective effects were exerted by AR overexpression in A375 cells treated with Dabrafenib individually and in combination with the MEK inhibitor Trametinib (TRA) (Fig. 2E).

Altogether, the findings indicate that AR overexpression in melanoma cells effectively counteracts growth suppression by BRAF inhibition.

### 3. Increased AR expression perturbs the transcriptional response of melanoma cells to BRAFi

For mechanistic insights, we undertook a global transcriptomic analysis of the three melanoma cell lines tested above. A large fraction of genes was similarly modulated in control and AR overexpressing cells at 48 hrs of Dabrafenib treatment (Fig. 3A, Suppl. Table 2). Gene families related to cell cycle and DNA replication were commonly downmodulated, consistent with the decreased rate of proliferation that also occurred with AR overexpressing cells at early times of Dabrafenib exposure (Fig. 3B, Suppl. Table 2). By contrast, the mitochondrial pro-apoptotic pathway genes were upregulated by Dabrafenib treatment to a much greater extent in control than AR overexpressing cells, consistent with the differential pro-apoptotic effects (Fig. 3B, Suppl. Table 2). Gene families related to pro-inflammatory signaling pathways (interferon α/ß and TNF-α) were significantly induced by Dabrafenib treatment selectively in control cells. Conversely, genes of the EGFR and TGF-ß pathways involved in melanoma progression and targeted drug resistance ^15,19^ were paradoxically induced by Dabrafenib treatment of AR overexpressing cells, to a much greater extent than in control cells (Fig. 3B, Suppl. Table 2). The findings were expanded by Gene Set Enrichment Analysis (GSEA), showing that signatures of interferon α and γ response were highly induced by Dabrafenib treatment of control cells, with a strong difference in these versus AR overexpressing cells (Fig. 3C, D). Importantly, an antigen presentation gene signature encompassing many major histocompatibility class I (MHC I) genes was also highly enriched in the Dabrafenib-treated control cells, with a profound difference relative to AR overexpressing cells (Fig. 3E).

Thus, increased AR expression in melanoma cells subverts the transcriptional response of melanoma cells to BRAF inhibition, with suppression of pro-apoptotic, immunomodulatory, and antigen presentation pathways and enhancement of pathways implicated in tumor progression and BRAFi resistance.

### 4. Increased AR expression elicits transcriptional changes of clinical significance found in BRAFi resistant subpopulations

To identify genes or sets of genes that are permanently modulated by increased AR expression and may account for their long-term BRAFi resistance, we compared the transcriptional profiles of the three melanoma cell lines plus/minus AR overexpression under basal conditions. Next to the *AR* gene itself, *SERPINE1*, an established TGF-ß target with pro-tumorigenic functions ^20,21^, was the single most upregulated gene in all three AR overexpressing melanoma cells (Fig. 4A, Suppl. Table 3). Together with a hallmark of AR response, the gene set enrichment analysis (GSEA) ^22^ showed a strong positive association of the profiles of AR overexpressing cells with predefined gene signatures related to cell proliferation (E2F), epithelial-mesenchymal transition (EMT) and undifferentiated and neural crest melanoma cells (UNDIF and UNDIF-NC) previously connected with BRAFi resistance ^23^ (Fig. 4B, Suppl. Table 3). Other gene signatures implicated in BRAFi resistance, specifically EGFR and TGF-ß signaling ^19^, were also positively associated with transcriptional changes elicited by AR overexpression (Fig. 4B, C; Suppl. Table 3).

**FIGURE 4:**
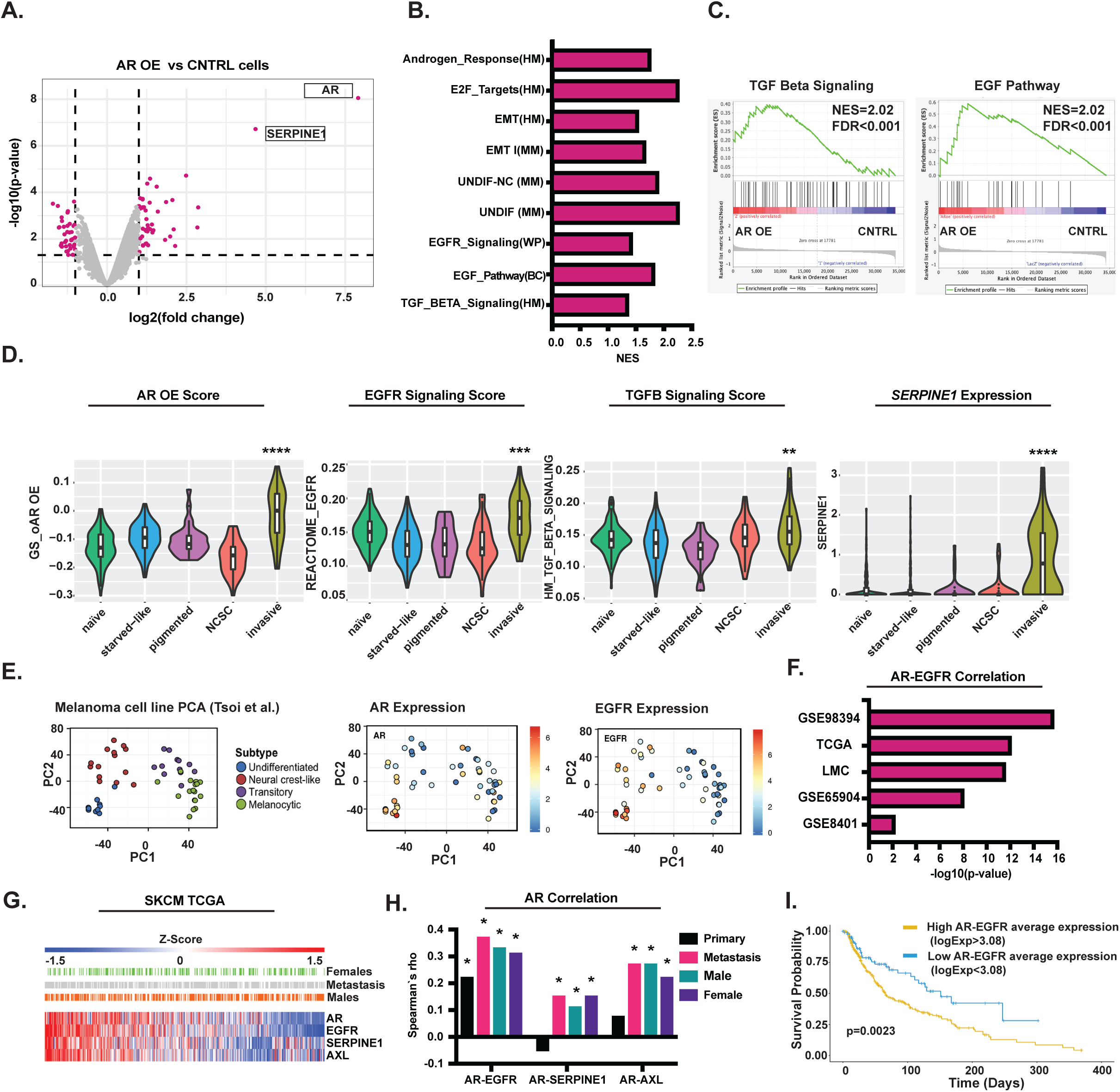
Increased AR expression elicits transcriptional changes of clinical significance found in BRAFi resistant subpopulations. A) Transcriptional changes elicited in melanoma cells by AR overexpression. Volcano plots of similarly modulated genes in A375, M14, and WM9 melanoma cells stably infected with AR versus LacZ expressing (control) lentiviruses under control conditions (w.o. Dabrafenib treatment). Plotting of differentially expressed genes is as in Fig. 3A. Coloured dots (magenta) correspond to genes with log_2_(fold change) threshold of -1 and 1 and p-value<0.05. The complete list of differentially expressed genes is provided in Suppl. Table 3. B) Gene Set Enrichment Analysis (GSEA) of transcriptional profiles of AR overexpressing versus control A375, M14, and WM9 melanoma cells as in the previous panel using a predefined set of gene signatures as obtained from the Hallmark gene set collection (HM) ^42^, wikipathways (WP) ^44^, biocarta (BC) and melanoma-specific studies (MM) ^23,37^ UNDIF-NC = undifferentiated, neural crest. Shown is a list of selected gene signatures with normalized enrichment score (NES) in profiles of AR overexpressing versus control melanoma cells. A more exhaustive list of signature genes is provided in Suppl. Table 3. C) GSEA and plot distribution of gene signatures related to EGF (biocarta) and TGF-ß (Hallmark) signaling. Genes are ranked by signal-to-noise ratio in AR-overexpressing versus control melanoma cells; the position of individual genes is indicated by black vertical bars; the enrichment pattern is in green. D) Scores of *AR* overexpression, EGFR, TGFß gene signatures activity, and *SERPINE1* expression in cell subpopulations identified by single cell RNA-seq analysis of a PDX model of melanoma BRAFi-resistance ^17^. A gene signature of 19 upregulated and 39 downregulated genes (absolute FC>1, p-value<0.01) in the AR overexpressing versus control melanoma cells was established (for a list of genes see Suppl. Table 3). The signature was used to calculate scores of AR activity, using AUCell ^40^, in the scRNA-seq profiles of previously defined populations of the drug-naive melanoma cells and BRAFi-induced starved-like (SMC), pigmented, invasive and neural crest-like subpopulations ^17^. Similar score calculations were performed with the Reactome EGFR signaling pathway ^45^ and the hallmark gene set for TGF-ß activity signatures ^42^ and single gene *SERPINE1* expression levels. Violin plots show individual cell score distribution within each cell population. The significance of differences in mean score values between invasive versus naïve cell populations (box plots) was calculated by Welch’s *t*-test ^41^. **, p<0.01; ***, p<0.001; ****, p<0.0001 E) Levels of AR and EGRF expression in multiple melanoma cell lines previously clustered according to multiple differentiation trajectories ^23^. Shown is the Principal Component Analysis (PCA) of expression profiles of individual melanoma cell lines (dots) and corresponding subtypes, together with overlapping color-coded indication of *AR* and *EGFR* mRNA levels, as retrieved from http://systems.crump.ucla.edu/dediff/. F) Positive correlation between AR and EGFR expression calculated from transcriptomic profiles of the indicated studies of melanoma clinical cohorts: GSE98394 (n=51 primary melanomas); TCGA (n=472 primary melanomas and melanoma metastases); LMC (n=703 primary melanomas); GSE65904 (n=214 melanoma metastases); GSE8401 (n=83 primary melanomas and melanoma metastases from xenograft models). Shown is the –log_10_ (p-value) of the correlation between AR and EGFR expression, as calculated using the *corrplot* v0.92 package with the Spearman’s correlation method. G) Heatmap of Z-score values for *AR, EGFR, SERPINE1* and *AXL* expression from RNA-seq profiles of 472 melanoma samples from the TCGA project (TCGA Firehose Legacy, February 2022). Z-scores were obtained by median-centering log_2_ (expression values) and dividing them by standard deviation. Shown are score values for each individual tumor, with a corresponding indication of patients’ sex, and whether they are from metastatic lesions. H) Correlation between *AR* an*d EGFR, SERPINE1* and *AXL* expression levels calculated from the TCGA melanoma cohort as in the previous panel, using the *corrplot* 0.92 package and the Spearman’s correlation method. Spearman’s rho coefficients are reported, with asterisks representing statistical significance (p-value < 0.05). I) Kaplan-Meier curves of long-term overall survival of melanoma patients from the TCGA dataset. Patients were divided according to high (yellow bar) versus low (blue bar) average expression of AR and EGFR, as calculated using the optimal cutpoint for continuous variables (log_2_ (Expression value) = 3.08), obtained from the maximally selected rank statistics from the *maxstat* R package.

A recent study of the BRAFi response at the single-cell level in mouse Patients Derived Xenografts (PDXs) pointed to a transition of drug-naive melanoma cells to a BRAFi-induced starved-like (SMC) subpopulation branching out to three phenotypes ^17^. By probing into the profiles of these distinct subpopulations, we found a highly enriched AR signature score in a specific BRAFi-tolerant subpopulation with elevated AXL expression and invasive features ^17^ (Fig. 4D). This same population was also found to have a positive enrichment score for the EGFR and TGF-ß gene signatures as well as *SERPINE1* expression (Fig. 4D).

These findings were extended by analyzing the composite transcription profiles of human melanoma cell lines that cluster into four main groups along a two-dimensional differentiation trajectory ^23^. Expression levels of the *AR* gene itself were positively associated with those of the *EGFR* gene in the most undifferentiated *AXL*-positive group connected with the targeted drug resistance ^23^ (Fig. 4E).

To assess the clinical significance of the results, we analyzed the transcriptomic profiles of melanoma cohorts, finding a strong positive correlation between expression levels of the *AR* and *EGFR* genes in multiple data sets (Fig. 4F). In the TCGA repository, we stratified lesions according to expression scores of the *AR, EGFR, SERPINE1*, and *AXL* genes (Fig. 4G). *AR* expression was positively associated with *EGFR* in both primary and metastatic melanoma lesions from male as well as female patients (Fig. 4H). *AR* expression also positively correlated with *SERPINE1* and *AXL* levels in the metastatic but not primary lesions, in keeping with the complex role played by these genes in melanoma progression. Lesions were subdivided according to optimal cutoff levels of average *AR* and *EGFR* expression, with patients with tumors with higher average expression levels having significantly lower survival than those with negative ones (log-rank test, p = 0.0026) (Fig. 4I). The findings remained significant after correcting for age, sex, and primary or metastatic status (multivariate Cox regression, p = 0.0073).

Thus, increased AR expression in melanoma cells elicits changes found in a BRAFi tolerant subpopulation and enhanced *EGFR* and *SERPINE1* expression of likely clinical significance.

### 5. Targeting AR overcomes BRAFi resistance

The above results suggested that AR is a positive determinant of melanoma progression and BRAFi resistance, which may be used for therapeutic targeting. Consistent with the transcriptomic results, RT-qPCR and immunoblot analysis showed a marked increase in *SERPINE1* (PAI-1) and *EGFR* levels in all tested melanoma cell lines upon AR overexpression (Fig. 5A, B).

**FIGURE 5:**
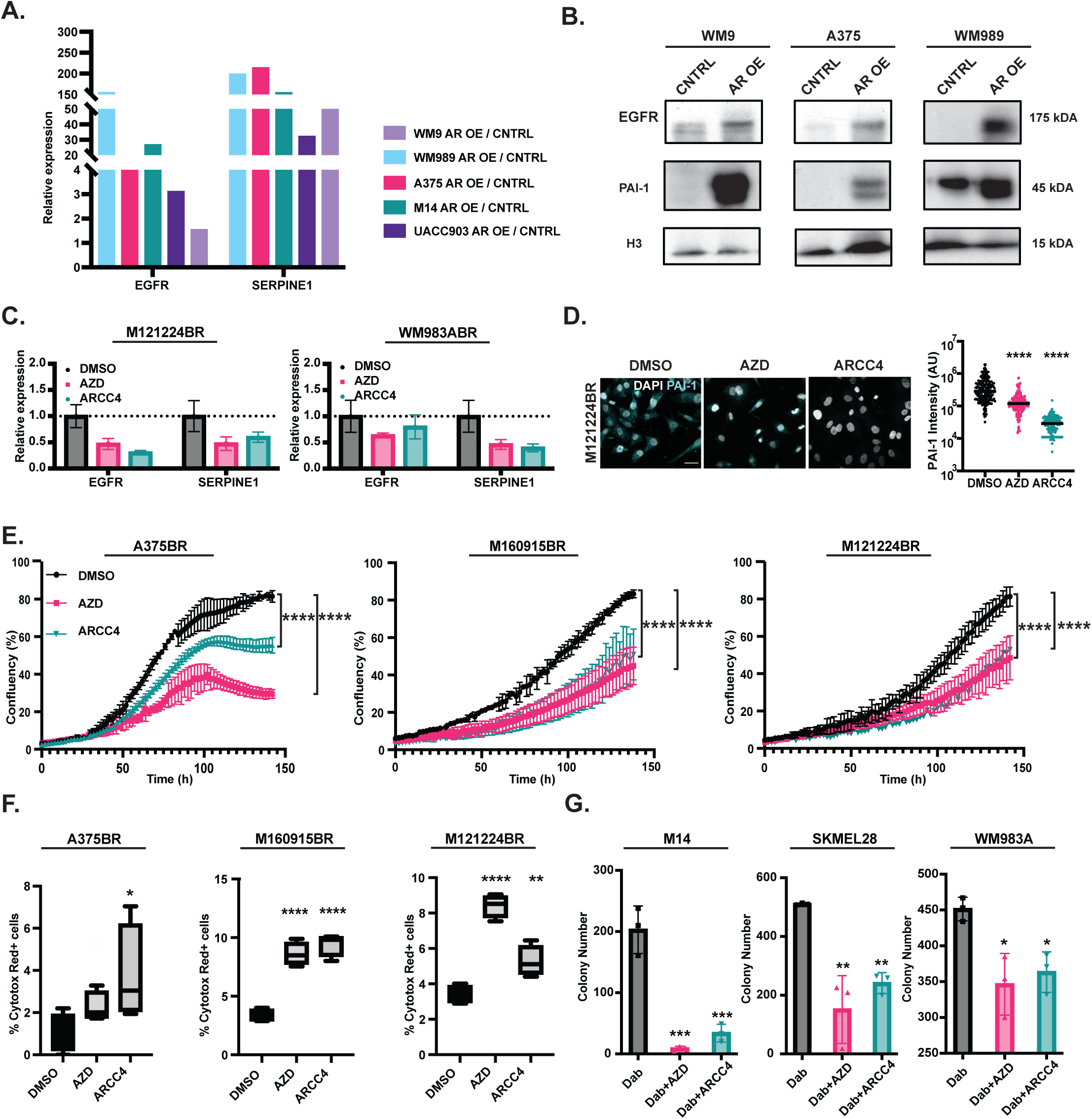
Targeting AR overcomes BRAFi resistance. A, B) Expression of the *EGFR* and *SERPINE1* genes in multiple melanoma cell lines plus/minus AR overexpression. The indicated melanoma cells stably infected with an AR overexpressing lentivirus versus LacZ expressing control (the same cells as in Fig. 4) were analyzed for expression of the *EGFR* and *SERPINE1* genes by RT-qPCR, with *RPLP0* for internal normalization (A) or immunoblotting with Histone H3 as an equal loading control (B). C) RT-qPCR analysis of *EGFR* and *SERPINE1* expression in two different melanoma cell lines (M121224BR and WM983ABR) treated with the AR inhibitors. Cells propagated in the presence of Dabrafenib as in Fig. 1 were treated with AZD3514 (10uM) or ARCC4 (1uM) versus DMSO control for 48 hours. Data are represented as the relative expression changes using *RPLP0* for internal normalization. D) Immunofluorescence analysis of melanoma cells (M121224BR) treated with the AR inhibitor AZD3514 or ARCC4 or DMSO control for 48 hours as in the previous panel with anti-PAI-1 antibodies with DAPI staining for nuclei visualization. Shown are representative images and quantification of the PAI-1signal intensity in arbitrary units (AU) per cell (dots) together with a mean, examining >100 cells per sample, unpaired *t*-test, **** p<0.0001. Color scale: grey, DAPI; cyan, PAI-1. Scale bar: 40μm. Immunofluorescence analysis of AR expression in parallel cultures is shown in Suppl. Fig. 4. E) Proliferation live-cell imaging assays (IncuCyte) of the indicated BRAFi resistant cells treated with AR inhibitors or DMSO as in the previous panels. Cells were plated in triplicate wells in 96-well plates, followed by cell density measurements (four images per well every 4 h for 128 h). cultures, n = 3; Pearson r correlation test. ****, p< 0.0001. F) Cell death as detected by live-cell staining (IncuCyte, Cytotox Red) of the same cultures as in the previous panel at 72 hours of treatment with the AR inhibitors versus DMSO control. Four images per well cultures, n (cultures) = 3; unpaired *t*-test, *, p<0.05; **, p< 0.01; ****, p<0.0001. G) Clonogenicity assays of three different drug-naive melanoma cell lines treated with Dabrafenib (0.5uM) individually or in combination with AZD3514 (10uM) or ARCC4 (1uM). Cells were plated in triplicates at clonal density (5000 cells / 6 cm dish) followed by 2 weeks of cultivation. Macroscopically detectable colonies were counted after crystal violet staining. n(dishes)=3, unpaired *t*-test, *, p<0.05; **, p< 0.01; ***, p<0.001.

To assess whether targeting of AR in BRAFi resistant melanoma cells elicits the converse effects, cells were treated with two different AR inhibitors, one suppressing AR activity through both ligand competitive and non-competitive mechanisms ^24^, and the other causing PROTAC-mediated degradation ^25^. Immunofluorescence and RT-qPCR analysis showed that treatment with both inhibitors caused effective loss of AR expression, which was paralleled by a decrease of *SERPINE1* (PAI-1) as well as EGFR expression (Fig. 5C, D, Suppl. Fig. 4).

The findings are of functional significance, as live-cell imaging assay showed that treatment with either AR inhibitors blunted melanoma cell proliferation and, at the same time, induced cell death (Fig. 5E, F). To assess whether inhibition of AR activity could also prevent the emergence of BRAFi resistance, drug-naive melanoma cells were cultured in the presence of Dabrafenib alone or in combination with the AR inhibitors. Consistent with previous studies ^15,26^, a large number of BRAFi-resistant colonies emerged in cultures of parental melanoma cells treated with the BRAFi alone, which was significantly reduced in cultures concomitantly treated with the AR inhibitors (Fig. 5G).

The studies were extended to an orthotopic model of melanoma development based on intradermal Matrigel injection of cells into immunodeficient mice. BRAFi-resistant A375 cells were treated with AZD3514 (10µM) versus DMSO vehicle alone 24 hours prior to injection. As shown in Fig. 6A, B, this single exposure to the AR inhibitor was sufficient to perturb the tumorigenicity of cells, which formed lesions with significantly decreased cell density and proliferation relative to controls.

**FIGURE 6:**
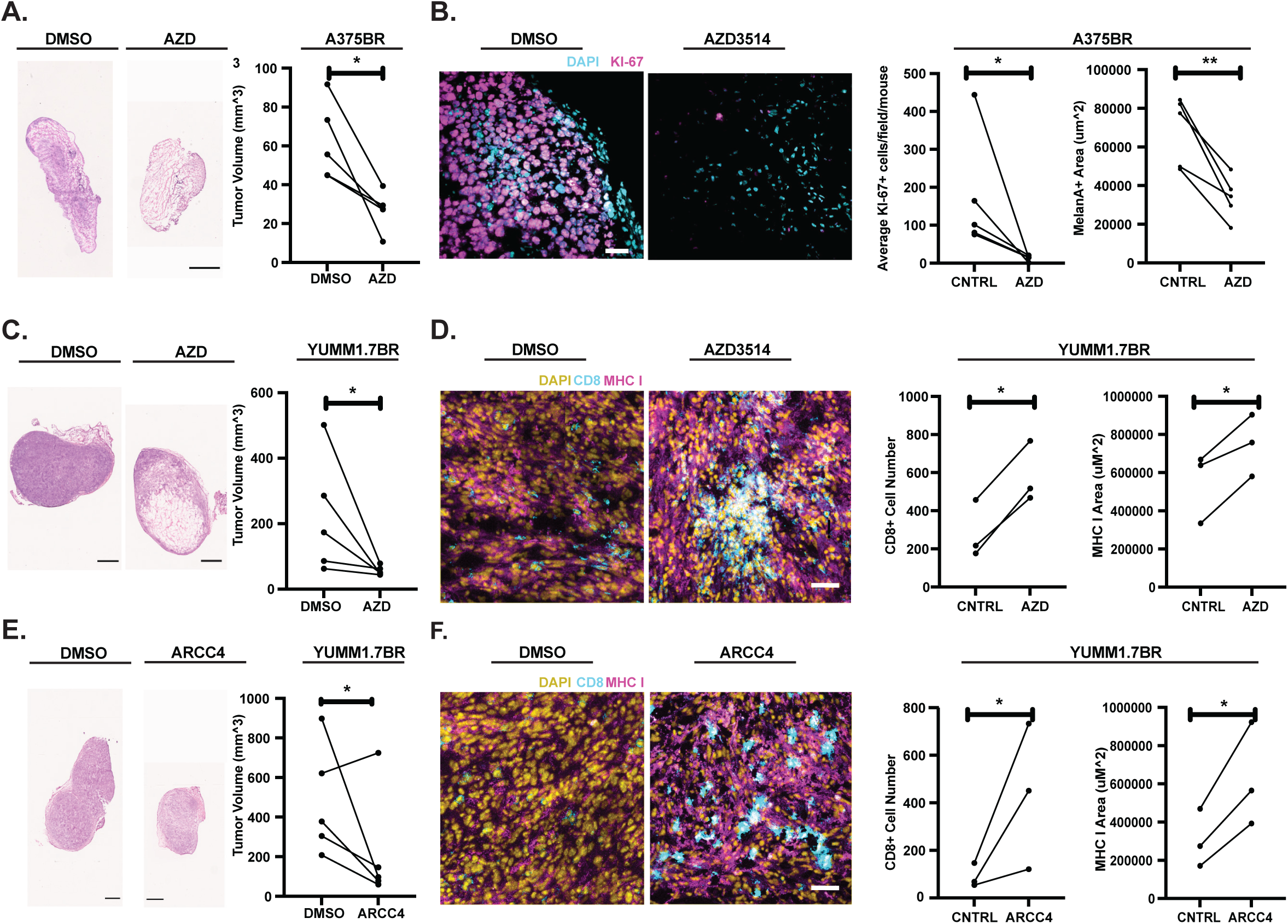
AR inhibition suppresses tumorigenicity of BRAFi-resistant melanoma cells. A) A375 cells selected for BRAFi resistance (A375BR) were subjected to a single treatment with the AZD3514 (10µM) versus DMSO control followed, 24 hours later, by parallel intradermal injections into 5 male NSG mice. Tumors were retrieved two weeks later. Shown are representative images of H&E stained lesions and quantification of tumor volume by caliper. n (mice) = 5, paired *t*-test, *, p<0.05. Scale Bar: 1 mm. B) Excised tumors, as in the previous panel, were analyzed by immunofluorescence analysis with antibodies against the Ki67 proliferation marker. Shown are representative images and quantification of the Ki67 proliferative index (number Ki67+ cells per field, three fields per lesion). n (mice) = 5, paired *t*-test, *, p<0.05. Scale Bar: 40μm. C-F) Mouse YUMM1.7 melanoma cells selected for BRAFi resistance (YUMM1.7BR), by multistep cultivation in increasing concentrations of Dabrafenib as with the human cells, were subjected to a single treatment with AZD3514 (10µM) (C, D) or ARCC4 (1µM) (E, F) versus DMSO control followed, 24 hours later, by parallel intradermal injections into 5 male C57BL/6JRj mice. C, E): representative images of H&E stained lesions and tumor volume quantification by caliper. n (mice) = 5, paired *t*-test, *, p<0.05. H&E Scale Bar: 1mm. D, F): double immunofluorescence analysis of excised tumors with antibodies against MHC I and the CD8+ T cell marker. Shown are representative images and quantification of MHC I + area and the number of CD8+ T cells per field (8 fields per lesion). n (mice) = 3, paired *t*-test, *p<0.05. Color scale: yellow, DAPI; magenta, PE-conjugated anti-MHC I antibody (MHC I-PE); cyan, anti-CD8+ antibody. Scale Bar: 40μm.

An important interconnection has been established in melanoma cells between the acquisition of BRAFi resistance and reduced sensitivity to the immune surveillance ^13^. The work was extended to a syngeneic mouse model, whereby BRAFi-resistant mouse melanoma cells (YUMM1.7) were pretreated with two different AR inhibitors, AZD3514 (10µM) or ARCC4 (1 µM), versus DMSO were injected into immunocompetent mice. Melanoma cells pretreated with both AR inhibitors produced tumors of significantly smaller size than controls with strongly increased MHC I surface expression and improved CD8+ T cell infiltration (Fig. 6C-F, Suppl. Fig. 5).

Hence, pharmacological inhibition of AR activity in BRAFi-resistant cells provides a tool to effectively suppress *EGFR* and *SERPINE1* expression, proliferation, and tumorigenicity with a concomitant enhancement of immune cell recognition.

## Discussion

Resilience to cancer therapy remains a major challenge even with improved approaches ^27^. In concert with modulation of the microenvironment, the resistance of cancer cells to targeted therapies can result from two mechanisms: i) an intrinsic adaptive response, with the expansion of pre-existing cell populations; ii) acquired resistance, through *de novo* genetic/epigenetic events ^27^. The adaptive response, which can be very rapid, is the result of compensatory feedback mechanisms of therapeutic interest ^28^. Drivers of adaptive responses are typically involved in regulatory circuits of both normal and cancer cells and can be most effectively targeted in the adjuvant therapy ^27^. Our combined findings indicate that the AR is one such driver as a key determinant of the adaptive response of melanoma cells to targeted therapy, which may be used to prevent or delay resistance.

The initial sensitivity of melanomas with activating BRAF mutations to BRAF inhibitors can be overcome by several mechanisms, including the compensatory upregulation of the EGFR tyrosine kinase coupled with downmodulation of the MITF and SOX10 transcription factors ^15,19,28,29^. In contrast to the negative role played by these transcription factors, we have found that AR is a positive determinant of BRAFi resistance and EGFR expression. We previously showed that basal AR activity is required for sustained proliferation and tumorigenesis of melanoma cells, with AR functioning as a bridge between RNA-Pol II and DNA repair proteins and ensuring the continuous DNA repair process associated with gene transcription ^9^. The markedly increased AR expression that is already occurring at early times of BRAFi and MEKi exposure suggested that this molecule can fulfill a second distinct function in melanoma cells as part of an adaptive mechanism leading to targeted drug resistance. In fact, persistently increased AR expression was by itself sufficient to render cells resistant to BRAFi-induced growth suppression and apoptosis, modulating different sets of genes from those affected by *AR* gene silencing ^9^ (see Suppl. Fig. 6 for a comparison).

Elevated AR expression in melanoma cells did not block but rather subverted the transcriptional response of melanoma cells to BRAF inhibition. Underlying the different sensitivity, apoptosis-related genes were induced by BRAFi treatment to a much greater extent in control than AR overexpressing cells. Efficacy of BRAF inhibitors depends on triggering a cancer cell death program associated with an impact on the tumor immune microenvironment ^30^. Gene signatures related to interferon signaling, inflammation, and antigen presentation, which can enhance immune stimulation and response to checkpoint inhibitors ^31^, were all induced by BRAFi treatment of control but not AR-overexpressing cells. A cross-connection has been established between BRAFi resistance and poor response to immune checkpoint control that does not depend on selection by the immune system and is a cancer cell-instructed ^13^, in which increased AR expression may be involved. Consistent with this possibility, in a syngeneic mouse model with BRAFi-resistant melanoma cells, low MHC I cell surface antigens expression, which has been linked with poor immune response ^32^, was strongly enhanced by treatment with AR inhibitors in parallel with CD8+ T cell infiltration.

Besides suppressing induction of pro-apoptotic and immunomodulatory genes, elevated AR expression resulted in a paradoxical upregulation by BRAFi treatment of gene families connected with BRAFi resistance, specifically EGFR and TGF-ß related ^19^. Gene signatures of two pathways were also induced by AR overexpression in melanoma cells under basal conditions, with an expression of the *EGFR* gene itself being consistently upregulated. Increased *EGFR* expression was previously connected with TGF-ß activation, with the two inducing cellular senescence of melanoma cells while, in the presence of BRAFi, conferring a growth advantage ^19^. The mechanism underlying this dichotomy remains to be established, and an interesting possibility is that AR is involved.

Among TGF-ß responsive genes, *SERPINE1* was prominently induced by increased AR expression. The gene codes for a secreted serine proteinase inhibitor of the SERPIN family (PAI-1), which exerts complex functions resulting from its binding to several cell surface proteins, promoting tumor development through effects on both cancer cells and the tumor microenvironment ^33^. It has been recently shown that elevated *SERPINE1* expression in melanoma cells is associated with a bad prognosis and poor response to immune checkpoint inhibitors ^20^.

The positive connection between *AR* and *EGFR* and *SERPINE1* expression was validated by single cell analysis of melanoma cells in a PDX model of BRAFi response: increased *AR* and *EGFR* gene signatures were coincidental with elevated *SERPINE1* expression in a specific BRAFi tolerant subpopulation characterized by high AXL expression and invasive features ^17^. Similarly, in a study on the heterogeneity of melanoma cell lines and tumors, *AR* expression was found to cluster together with *EGFR* and *SERPINE1* in an undifferentiated AXL positive subgroup connected with targeted drug resistance ^23^. The positive association of *AR* with *EGFR, SERPINE1*, and *AXL* expression was confirmed in a large patient cohort, irrespective of sex and primary versus metastatic lesions, with poor survival with tumors with elevated *AR*-*EGFR* levels.

AR has been intensely studied as a driver of metastatic prostate cancer, with resistance to AR-targeting approaches resulting from various mechanisms, including increased *AR* expression ^34^. *AR* upregulation is a point of convergence of multiple mechanisms ^18^, with transcription factors like CREB1 and Foxo3a that we have found to be positively associated with *AR* transcription also in our system. Overall, genetic and epigenetic changes of *AR* resistance are less likely to occur in melanoma, in which other genes drive the disease. Inhibitors targeting AR activity and expression could be employed as co-adjuvants to prevent/delay targeted drug resistance and, as we have shown, suppress tumorigenicity of BRAFi-resistant cells while at the same time inducing CD8+ T cells infiltration. Given the connection between BRAFi resistance and poor immune response ^13^, as well as the intrinsic role of AR activation in dampening the T cell activity ^12,35^, AR targeting may be beneficial in the treatment regimens with immune checkpoint inhibitors.

## Materials and methods

### Cell Culture

A full list of different melanoma cell lines and primary melanoma cells is provided in Supplementary Table S4. Early passage primary melanoma cell cultures (M160915 and M121224) were established from discarded melanoma tissue samples by the University Research Priority Program (URPP) Live Cell Biobank (University of Zurich) following institutional requirements. WM115, WM9, WM983A, WM989, UACC903, and UACC903BR melanoma cells were a gift from Dr. Meenhard Herlyn (The Wistar Institute, US). The YUMM1.7 melanoma cell line ^36^ was provided by Dr. Ping-Chih Ho (UNIL). All melanoma cell lines and patient-derived primary melanoma cells were maintained in Roswell Park Memorial Institute (RPMI) medium (Thermo Fisher Scientific) supplemented with 10% (v/v) fetal bovine serum (Thermo Fisher Scientific). YUMM1.7 melanoma cells were maintained in Dulbecco’s Modified Eagle Medium (DMEM) (Thermo Fisher Scientific) supplemented with 10% (v/v) fetal bovine serum (Thermo Fisher Scientific). All cell lines were routinely tested for *Mycoplasma*. Cell morphology and growth characteristics were monitored throughout the study and compared with the previously published reports. No further authentication of these cell lines was performed.

### Cell manipulations and treatments

*Lentiviral particle productions and infections* were performed as described previously ^9^. Melanoma cells were transduced with *AR* overexpressing (a gift of Dr. Karl-Henning Kalland, Bergen University, Bergen, Norway) or LacZ expressing control lentiviruses for 6 hours. Two days post-infection cells were selected using 5 μg/ml of Blasticidin for 6 days. RNA or protein samples were collected 7 days after infection.

*BRAF resistant (BR) cell lines* were established from the parental (P) cells (A375, M160915, M121224, and WM983A) by continuous culturing in Dabrafenib for a period of 4 weeks, with weekly multistep increases in concentration from 0.5 to 3 µM. Resistant cells were thereafter continuously cultured in the presence of 3 µM Dabrafenib.

For *short-term in vitro experiments with various chemical inhibitors*, 24 hours post-seeding cells were treated with the following compounds at the indicated concentrations: Dabrafenib (0. 5 µM), PLX-4720 (0. 5 µM), Sorafenib (0. 5 µM), Trametinib (0.005 µM), Cobimetinib (0.005 µM), s31-201 (50 µM), Bay 7085 (10 µM), T55224 (20 µM), sr11302 (10 µM), SU6668 (10 µM), SKI-606 (10 µM), and CYT387 (10 µM). All inhibitors were purchased from SellectChem and were dissolved in DMSO according to the manufacturer’s instructions. DMSO was used as vehicle control. RNA was collected 48 hours post-treatment. For *AR inhibition*, 24 hours post-seeding, melanoma cells were treated with 10 µM AZD3514 (SellectChem) or with 1 µM ARCC4 (Tocris). AR inhibitors were dissolved in DMSO according to the manufacturer’s instructions. DMSO was used as vehicle control for all experiments.

*Cell proliferation / density assays* were carried out by measuring ATP production using the CellTiter-Glo luminescent assay (Promega) as per the manufacturer’s instructions. Dabrafenib dose-response curves and IC50 values were attained by fitting the curves to nonlinear regression with variable slope using GraphPad Prism.

For *clonogenicity assays*, cells were plated onto 60 mm dishes (10,000 cells/well; triplicate wells/condition) and treated the next day as indicated in the figure legends. Tissue culture medium was refreshed every 2-3 days. Cells were cultured for 7 days for *AR* overexpressing experiments and 14 days for experiments with AR inhibitors. Colonies were fixed with methanol fixed and stained with 1% crystal violet. The number of clones was counted using Fiji/ImageJ software.

For *IncuCyte cell proliferation and cell death assays*, 1000 melanoma cells per condition were seeded in triplicate in 96-wells plates. Drug treatments were applied 12 hours post-seeding, with cells allowed to proliferate for 5 days. Cell proliferation was monitored using the IncuCyte Zoom Live-Cell Imaging System (Essen Bioscience). Four independent images per well per condition were captured every 2 hours for 5 days. Cell confluence was determined with IncuCyte Zoom software. For cell death measurements, the IncuCyte Cytotox Red Reagent was added to the cells seeded and treated as described above and imaged according to the manufacturer’s instructions. Cytotox Red positive cells were quantified using the IncuCyte Zoom software.

### Immunofluorescence and immunohistochemistry staining

Immunofluorescences staining of tissue sections and cultured cells was carried out as described previously ^9^. In brief, paraffin-embedded sections were deparaffinized and rehydrated prior to a citrate-based buffer antigen retrieval. Frozen tissue sections (8 mm) or cultured cells were fixed in 4% paraformaldehyde (PFA) for 15 minutes at room temperature (RT). Samples were washed with PBS (3×5min) and permeabilized using 0.5% TritonX100 in PBS for 10 minutes. Samples were blocked using 2% bovine serum albumin in PBS for 1 hour at RT. Primary antibodies were diluted in a blocking buffer (PBS/2% bovine serum albumin) and were incubated overnight at 4°C. Following, samples were washed (PBS, 3×5min) and incubated with secondary donkey fluorescence conjugated secondary antibodies (Invitrogen) for 1 hour at RT. DAPI was used to counterstain nuclei. Slides were washed (PBS, 3×5min) and mounted using Fluoromount Mounting Medium (Sigma-Aldrich). Control staining without the primary antibodies was performed in each case to subtract background and set image acquisition parameters. A full list of primary and secondary antibodies and dilutions used for IF is provided in Supplementary Table 5.

Immunofluorescence images were acquired with a ZEISS LSM880 confocal microscope with 20X, 40X, or 63x oil immersion objectives or with a NanoZoomer S60 microscope with a 40X objective. ZEN Blue software was used for image acquisition. Fiji/ImageJ software was used for image processing and analysis. For image analysis, the images were stacked to maximal projections, and immunofluorescent channels were split. A binary mask was then created using a watershed function in the DAPI channel, allowing for the identification of individual nuclei. The mean grey value intensity of channels was measured and summed. The fluorescent intensities are indicated in arbitrary units. On average, >100 cells were analyzed for *in vitro* studies. For *in vivo* studies, five fields were imaged per tumor.

Immunohistochemical staining was performed by the laboratory of pathology in the Department of Biochemistry, UNIL, as previously described ^9^. Slides were scanned using a NanoZoomer S60 microscope with a 20X objective. Ndp.View2 and Fiji/ImageJ software were used for the acquisition and processing of images.

### Immunoblotting

Cells were lysed using boiling LDS buffer (2%SDS, 50□mM Tris/HCl (pH 7.4) supplemented with 1□mM PMSF, 1□mM Na3VO4, and 10□mM NaF. Total protein content was quantified with a BCA assay (Thermo Fisher Scientific). Equal amounts (20-50 µg) of proteins were subjected to 10% SDS–PAGE followed by immunoblot analysis. All membranes were sequentially probed with different antibodies as indicated in the figure legends. Super Signal West Pico PLUS Chemiluminescent Substrate (Thermo Fisher Scientific) was used for signal detection. Full details of antibodies used in this study are provided in Supplementary Table 5.

### Real-time quantitative PCR (RT-qPCR)

Total mRNA was extracted using TRIzol according to the manufacturer’s instructions, followed by cDNA synthesis using the RevertAid H Minus Reverse Transcriptase (Thermo Fisher Scientific). RT-qPCR was performed using SYBR Fast qPCR Master Mix (Kapa Biosystems) on a Light Cycler 480 (Roche). The relative quantification (RQ) and expression of each mRNA were calculated using the comparative Ct method. All samples were run in technical triplicates and were normalized to an endogenous control, *RPLP0*. A full list of primers used in the study is provided in Supplementary Table 5.

### Transcriptomics and bioinformatic analysis

A375, M14, and WM9 melanoma cells infected with an *AR* overexpression lentivirus versus LacZ expressing control virus were treated with Dabrafenib (0.5 µM) versus DMSO vehicle for 48 hours. Following, RNA was extracted using the Direct-zol RNA MiniPrep kit (Zymo Research) coupled with DNase treatment according to the manufacturer’s instructions. The RNA quality was first evaluated using Agilent 2100 Bioanalyzer® (Agilent Technologies, USA). Transcriptomic analysis was performed using Clariom™ D GeneChip array hybridization (Thermo Fisher Scientific). Single-strand cDNA preparation, labeling, and hybridization were performed in accordance with Affymetrix protocols at the iGE3 Genomics Platform, University of Geneva (Geneva, Switzerland). Data obtained (CELL files) were summarized using the RMA function in the R package oligo with background correction and quantile normalization. Gene IDs were mapped using the Chip-annotation package clariomdhumantranscriptcluster.db. The R package “limma” was used for gene differential expression analysis, followed by multiple testing correction by the Benjamini-Hochberg procedure. The cutoffs for the Dabrafenib treatment signatures (Dabrafenib CNTRL vs DMSO CNTRL and Dabrafenib AR OE vs AR OE CNTRL) were FC > 1.5, and adj-p < 0.01, yielding 360 up- and 360 down-regulated genes for CNTRL Dabrafenib-treated cells, and 199 up- and 344 down-regulated genes for AR OE Dabrafenib-treated cells. The cutoff for the AR OE signature (AR OE CNTRL vs DMSO CNTRL) was FC > 2.0, and p-value < 0.05, yielding 48 up- and 39 down-regulated genes. The data generated in this study have been deposited to the public functional genomics data repository GEO (Gene Expression Omnibus), NCBI with an accession number **GSE199405**.

*Gene ontology (GO) enrichment analysis* was performed on the differentially expressed genes with the fold change cutoff value of 2.0 using the Enrichr. Gene Ontology and Pathway Classification System to identify the enriched biological processes.

*Gene set enrichment analysis (GSEA)* for GeneChip microarray data was conducted using GSEA software using default parameters. Curated gene sets were obtained from various sources as also indicated in the legends for Figs. 3 and 4: i) the Molecular Signatures Database (MSigDB version 5.2, www.broadinstitute.org/gsea/msigdb/); ii) previously published melanoma-specific signatures ^17,23,37^. A list of enriched pathways is provided in Supplementary Tables S2,3.

For *AR overexpression score*, CELL files were summarized using the RMA function in the R package oligo with background correction and quantile normalization. Gene IDs were mapped using the Chip-annotation package clariomdhumantranscriptcluster.db and differential expression analysis was performed with limma, using the formula ∼treatment + cell_line. The treatment referred to the comparison DMSO versus AR OE.

*Signature score analysis of single cell RNA-seq profiles* was performed starting from single cell RNA-seq data (GEO # GSE116237) filtering for cells with more than 1000 gene counts and genes detected in more than 3 cells. Further filtering was omitted as it has already been done by the authors of the dataset ^17^. Ensembl IDs were mapped into gene symbols using biomaRt ^38^ and count data were summed together when multiple IDs mapped to the same symbol. Library normalization, log transformation and further downstream analysis were performed using Seurat v4 ^39^. Signature scores were calculated using AUCell ^40^ and significance between scores or individual gene expressions were calculated using Welch’s *t*- test ^41^. Gene sets were downloaded from the Molecular Signatures Database ^42^.

*Correlation analysis* between AR, EGFR, PAI-1, and AXL expression levels was calculated on 472 melanoma samples from the TCGA project (TCGA Firehose Legacy, February 2022) with the corrplot 0.92 package, using the Spearman’s correlation method.

*Survival analysis* was based on the melanoma TCGA dataset and calculation of the optimal cutpoint for continuous variables (log2Expression value = 3.08) from the maximally selected rank statistics from the ‘maxstat’ R package.

### *In Vivo* Studies

NOD *scid* gamma (NSG) mice, (NOD.Cg-Prkdcscid Il2rgtm1Wjl/SzJ; 6-8-week-old males), were obtained from the Jackson Laboratory. BRAFi resistant human A375 melanoma cells (A375BR) were pretreated with AZD3514 (10 μM) or DMSO control for 12 hrs prior to injection into mice. Cells (1 × 10^6^ per injection, in Matrigel (Corning), 70 μl) were injected intradermally in parallel into the left and right flanks of mice with 29-gauge syringes. Mice were sacrificed and Matrigel nodules were retrieved for tissue analysis 10 days after injection.

C57BL/6JRj mice (6-8-week-old males) were obtained from Jackson Laboratory. BRAFi resistant murine melanoma cells (YUMM1.7BR) were pretreated with AZD3514 (10 μM), ARCC4 (1 μM), or DMSO control for 12 hrs prior to injection. 2 × 10^6^ melanoma cells per condition were injected with Matrigel (Corning) (70 μl per injection) intradermally in parallel into the left and right side of mice with 29-gauge syringes. Mice were sacrificed and Matrigel nodules were retrieved 14 days after injection. Tumors were measured post- extraction using calipers. Tumor volumes were calculated using the formula (LxW^2^x0.5). Throughout the study, general humane endpoints were applied. All mice were housed in the animal facility of the University of Lausanne.

### Statistical analysis

All statistical tests were performed using GraphPad Prism 9 (GraphPad Software, Inc.). Data are shown as mean± SEM or mean ± SD, as indicated in the legends. Detailed information on the statistical methods applied for each experiment can be found in the corresponding figure legends. Statistical difference between two groups was determined using Student’s *t*-test unless otherwise mentioned. For comparisons among more than two groups, a one-way analysis of variance (ANOVA) followed by Bonferroni’s correction was used. For longitudinal data, Spearman’s correlation was used to infer significance between the experimental treatment arms.

For tumorigenicity assays, individual animal variability issue was minimized by contralateral injections in the same animals under control versus experimental conditions. No statistical method was used to predetermine sample size in animal experiments and no exclusion criteria were adopted for studies and sample collection. No exclusion criteria were adopted for animal studies or sample collection. No randomization was used, and the researchers involved in the study were not blinded during sample obtainment or data analysis.

### Study approvals

Pre- and post-treatment metastatic melanoma sections were obtained from the Live Cell Biobanks of the University Research Priority Program (URPP) “Translational Cancer Research” (Mitchell P. Levesque, University Hospital Zurich). All human samples were obtained from surplus melanoma material collected from de-identified patients who provided written, informed consent to participate in the research (BASEC-Nr 2017—00494). No access to sensitive information has been provided.

All animal studies were carried out according to Swiss guidelines for the use of laboratory animals, with protocols approved by the University of Lausanne animal care and use committee and the veterinary office of Canton Vaud (animal license No. 1854.4f/1854.5a).

## Acknowledgments

We are grateful to Meenhard Herlyn for gifting WM115, WM9, WM989, WM983A, UACC903, and UACC903BR melanoma cell lines used in the study. We thank Ping-Chih Ho for a gift of murine YUMM1.7 melanoma cells. We are grateful to Luigi Mazzeo, Jovan Isma, and Sandro Goruppi for stimulating discussions. We thank the URPP in Translational Cancer Research Biobank at the University of Zürich for access to the melanoma cell lines and patient samples. This work was supported by grants from the Swiss Cancer Research Foundation (project number: KFS-4709-02-2019) and the National Institutes of Health (R01AR078374; R01AR039190; content not necessarily representing the official views of NIH). The laboratory has also received funding from the European Union’s Horizon 2020 research and innovation program under the Marie Skłodowska-Curie grant agreement No 859860. GPD and MPL are members of the SKINTEGRITY.CH collaborative research program.

## Author contributions

AS, SG, TP, BT, and MM performed experiments and analyzed the results with GPD. MKY, PO, and GC performed the bioinformatic analysis, MPL provided the clinical samples and critical feedback. AS and GPD designed the study and wrote the manuscript.

## SUPPLEMENTAL INFORMATION SUPPLEMENTARY FIGURES

**Supplementary Fig. 1:** BRAFi treatment of melanoma cells results in increased AR expression.

**Supplementary Fig. 2:** Increased AR expression in clinical post-BRAFi/MEKi treatment melanoma samples.

**Supplementary Fig. 3:** BRAFi treatment induces expression of positive regulators of AR expression.

**Supplementary Fig. 4:** Decreased AR protein expression by treatment of melanoma cells with AR inhibitors.

**Supplementary Fig. 5:** Pharmacological AR targeting alters MHC I surface expression and CD8 T cell infiltration.

**Supplementary Fig. 6:** Comparative analysis of transcriptomic profiles of melanoma cells plus/minus AR gene silencing versus overexpression.

## SUPPLEMENTARY TABLES

**Supplementary Table 1:** Gene expression levels of selected transcription factors upon Dabrafenib treatment and Spearmans’s correlation scores from the TCGA.

**Supplementary Table 2:** List of differentially expressed genes and gene ontology terms enriched in the control and AR overexpressing cells plus/minus Dabrafenib treatment.

**Supplementary Table 3:** List of differentially expressed genes and GSEA terms enriched in AR overexpressing cells under basal conditions.

**Supplementary Table 4:** List of cell lines used in the study.

**Supplementary Table 5:** List of key reagents and resources used in the study.

