## Supplementary for "Androgen Receptor is a Determinant of Melanoma targeted drug resistance"

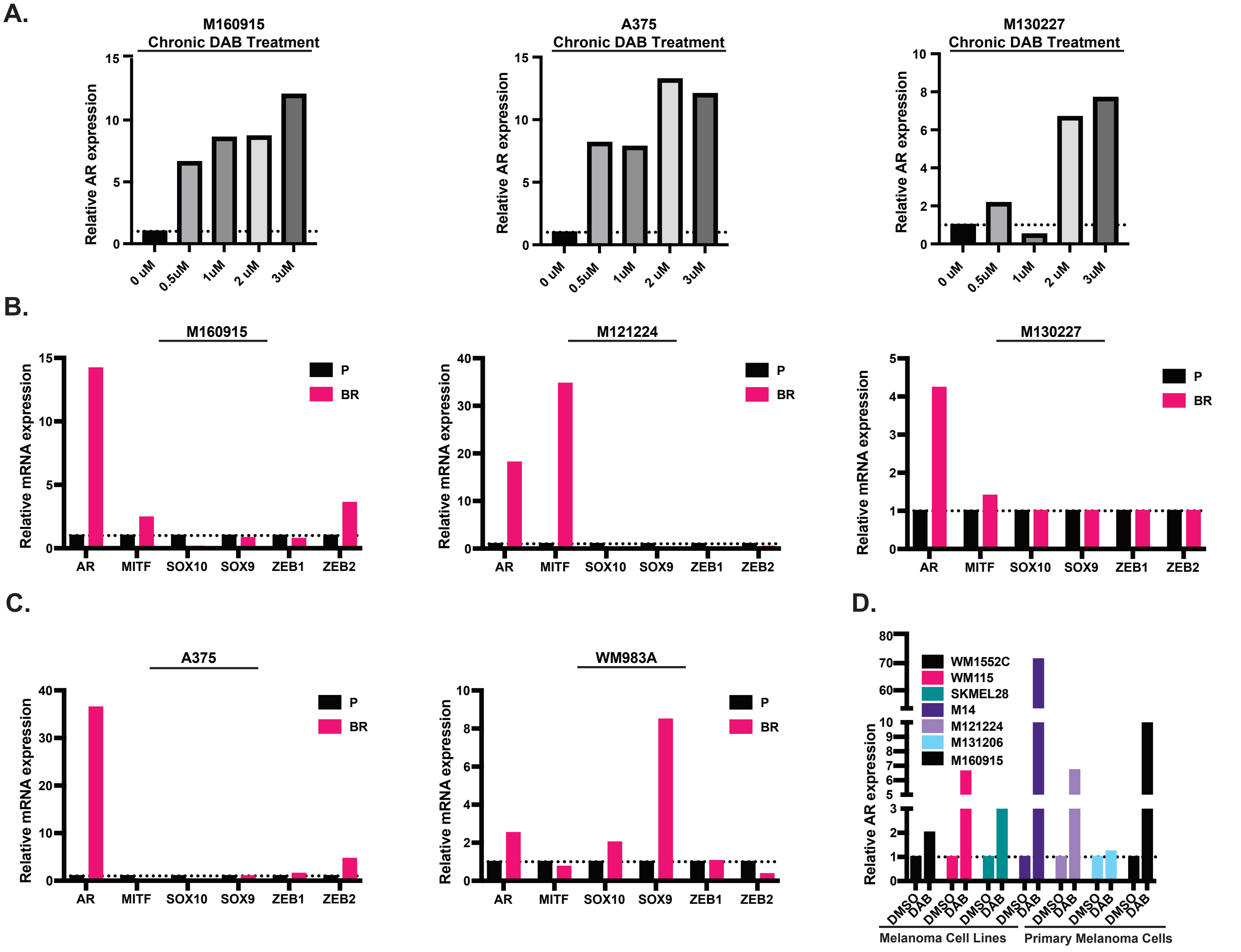


**Supplementary Fig. 1: BRAFi treatment of melanoma cells results in increased AR expression.**
A) *AR* expression in three additional human primary and established melanoma cell lines. Cells were cultured with multistep weekly increases of the BRAF inhibitor Dabrafenib (0.5, 1, 2, and 3 μM). Cells were collected at the end of each week of treatment and analyzed together with the untreated parental cells for levels of *AR* expression by RT-qPCR, with *RPLP0* for internal normalization. Related to Fig. 1A.
B-C) Relative expression of the indicated genes in BRAFi resistant (BR) primary (B) and established melanoma cell lines (C) versus parental cells (P). Shown are individual bar plots of the heatmap shown in Fig. 1D. RT-qPCR results are expressed as fold changes relative to untreated controls, after *RPLP0* normalization.
D) Relative *AR* expression in the indicated primary and established melanoma cells at 48 hours of treatment with BRAF inhibitor Dabrafenib (0.5 μM) versus DMSO control. RT-qPCR results are expressed as fold changes relative to untreated controls, after *RPLP0* normalization. Related to Fig. 1E.

**Supplementary Figure 1.**

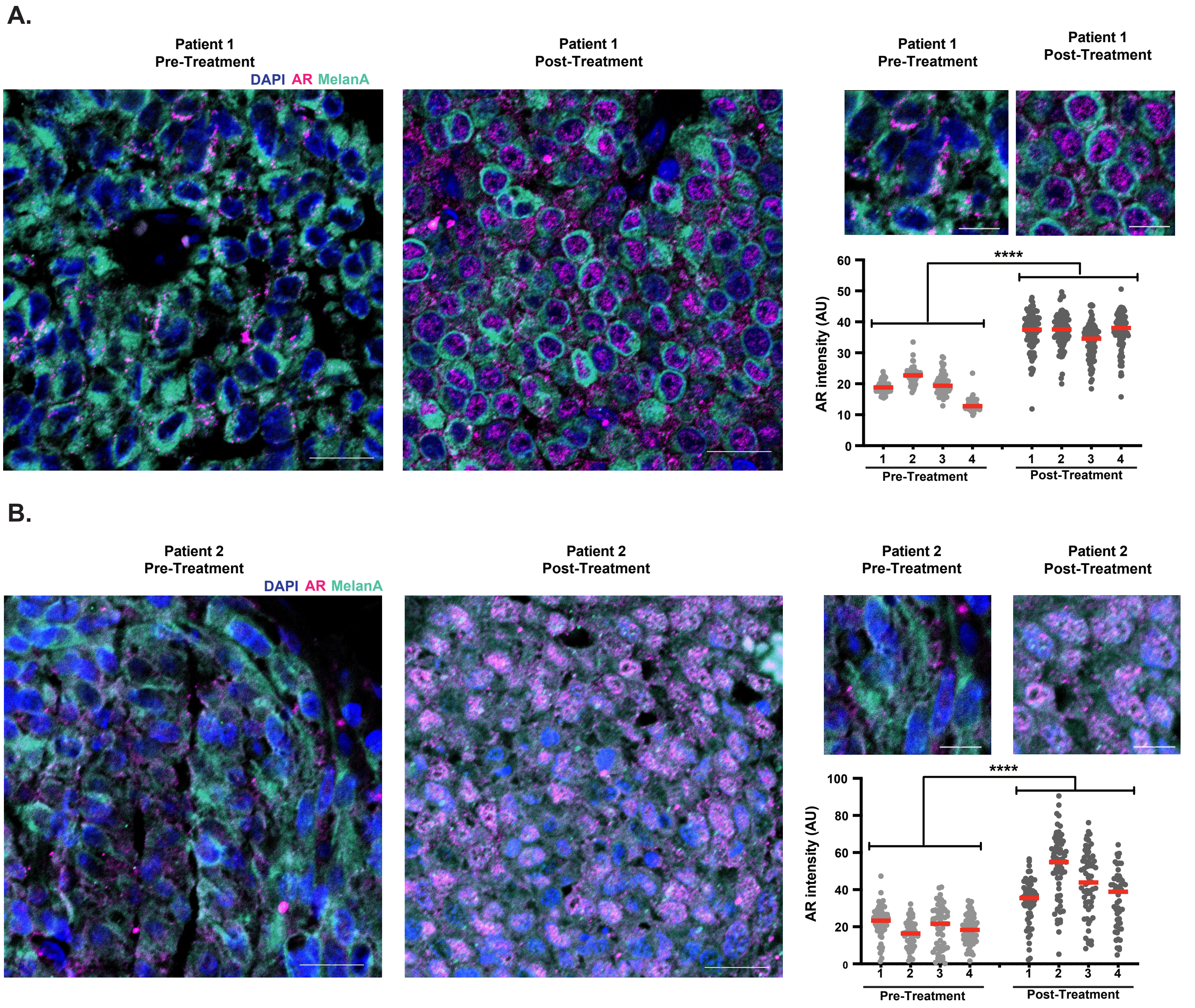


**Supplementary Fig. 2: Increased AR expression in clinical post-BRAFi/MEKi treatment melanoma samples.**

A-B) Immunofluorescence analysis of matched pre-and post- BRAFi/MEKi treatment samples (patients 1 (A) and 2 (B)). Shown are representative low- and high-magnification images of the areas used for single-cell AR expression quantification. Quantification of the AR signal intensity in arbitrary units (AU) per cell (dots) together with a mean, examining >50 cells per field (4 fields per lesion), paired *t*-test, **** $p<0.0001$. Color scale: blue, DAPI; cyan, MelanA; magenta, AR. Scale bar: 20 µm and 10 µm, respectively.

**Supplementary Figure 2.**

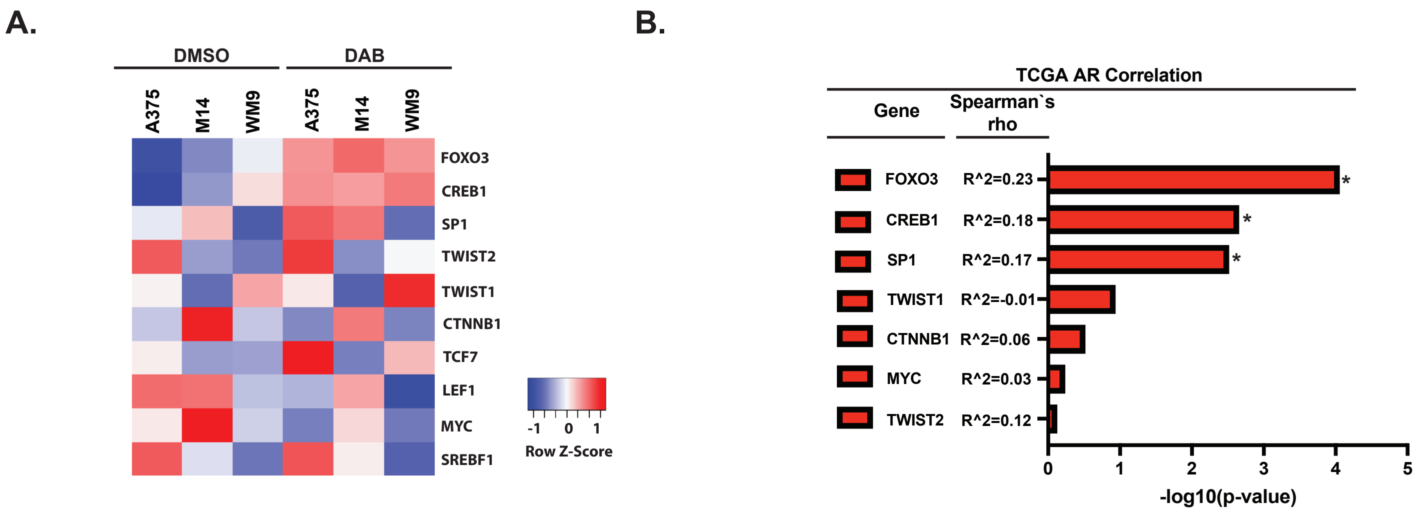


**Supplementary Fig. 3: BRAFi treatment induces expression of positive regulators of AR expression.** A) Heatmap of expression changes in the indicated transcription factors in three melanoma cell lines (A375, M14, and WM9) plus/minus Dabrafenib (48 hours), as obtained from global transcriptomic profiles (Suppl. Table 2). Z-scores were obtained by mean-centering log2 (expression values) and dividing them by standard deviation. Shown are z-scores for each individual experimental condition.
B) Correlation analysis between *AR* and indicated transcription factors calculated from the TCGA transcriptomic profiles (n=472 primary melanomas and melanoma metastases). Shown are the –log10 (p-value) (bar plots) and the Spearman`s rho coefficient of the correlation between AR and indicated transcription factor expression, as calculated using the corrplot v0.92 package with the Spearman`s correlation method.

**Supplementary Figure 3.**

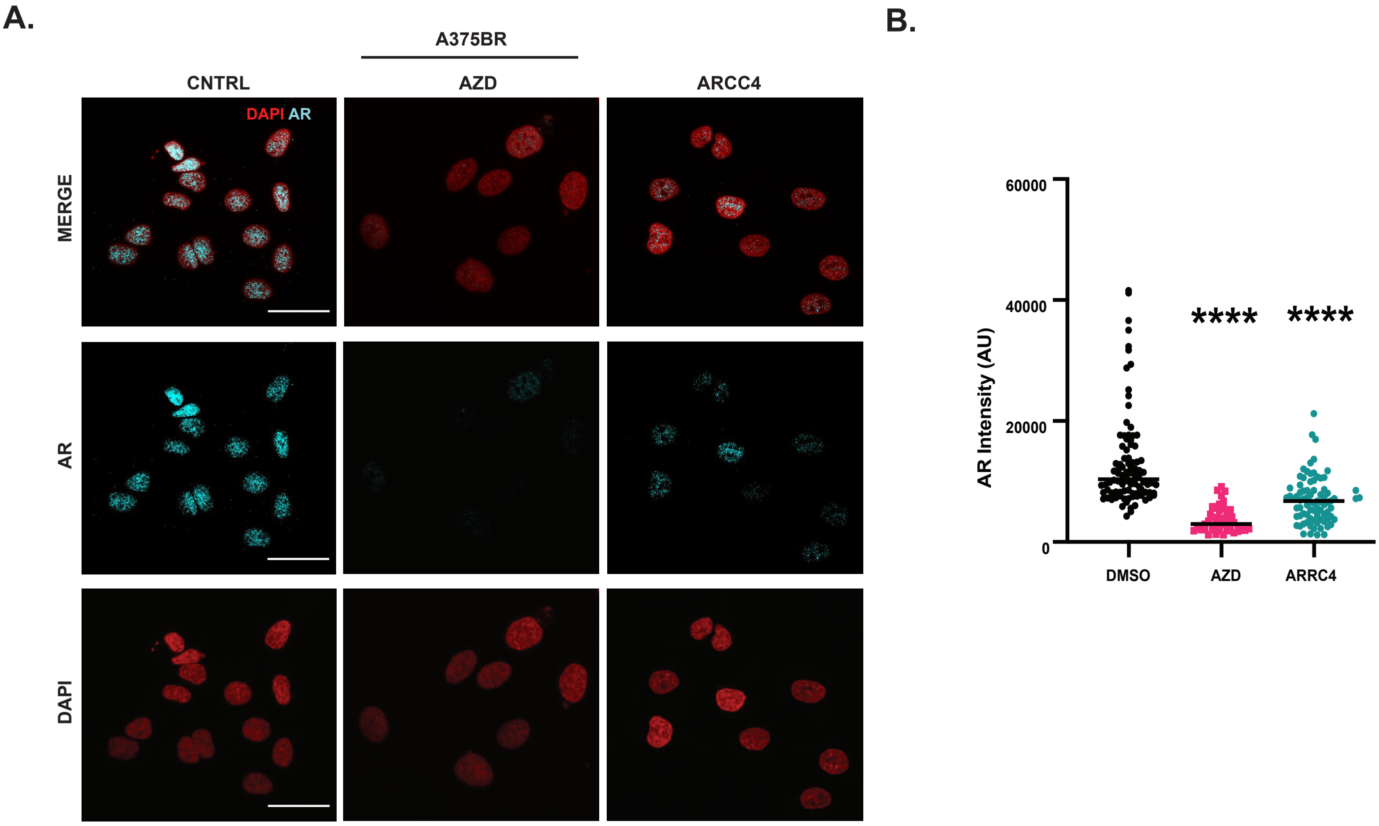


**Supplementary Fig. 4: Decreased AR protein expression by treatment of melanoma cells with AR inhibitors.**

A-B) Immunofluorescence analysis of BRAF inhibitor-resistant melanoma cells (A375BR) treated with the AR inhibitor AZD3514 (10µM) or ARCC4 (1µM) versus DMSO control for 48 hours with anti-AR antibodies with DAPI staining for nuclei visualization. Shown are representative images (A) and quantification (B) of AR signal intensity, as obtained from digitally acquired images and ImageJ analysis, in arbitrary units (AU) per individual cell (dots) together with a mean, examining >100 cells per sample, unpaired t-test, **** p<0.0001. Color scale: red, DAPI; cyan, AR. Scale bar: 40 µm. Related to Fig. 5D.

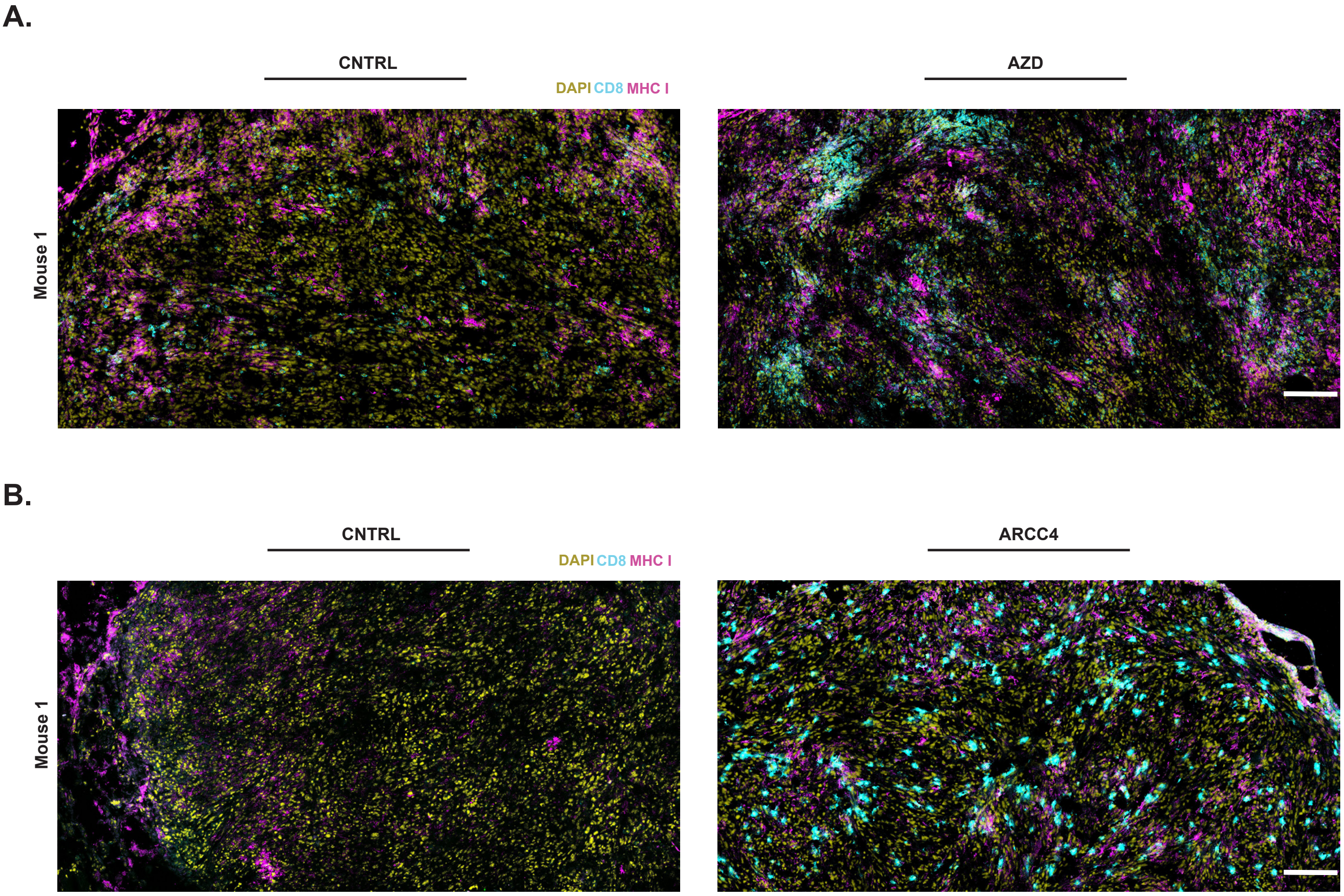


**Supplementary Fig. 5: Pharmacological AR targeting alters MHC I surface expression and CD8+ T cell infiltration.**

A-B) Immunofluorescence analysis of the excised YUMM1.7BR tumors subjected to a single treatment with AZD3514 (10 μM) (A) or ARCC4 (1 μM) (B), as described in Fig. 6. Shown are representative low magnification images of the MHC I + area and the CD8+ T cells. Color scale: yellow, DAPI; magenta, PE-conjugated anti-MHC I antibody (MHC I-PE); cyan, anti-CD8+ antibody. Scale Bar: 100 μm. Related to Fig. 6D, F.

Supplementary Figure 5.

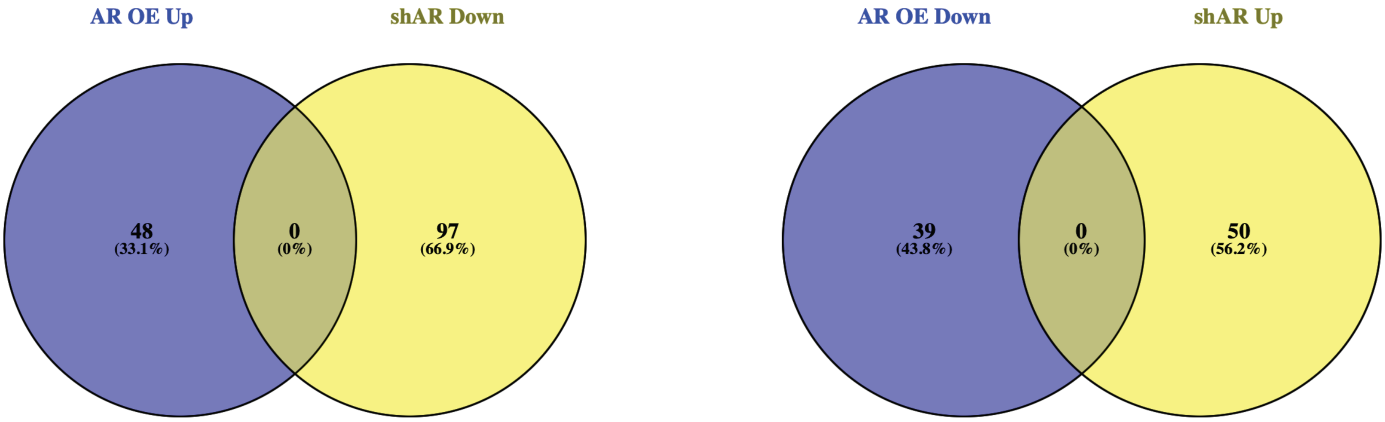


**Supplementary Fig. 6: Comparative analysis of transcriptomic profiles of melanoma cells plus/minus AR gene silencing versus overexpression.**
Venn diagrams illustrating common and differentially expressed genes in multiple melanoma cell lines plus/minus *AR* gene silencing as previously reported[9] versus those in melanoma cells plus/minus *AR* overexpression (Suppl. Table 3). Number of significantly modulated genes that were oppositely modulated in the two conditions (FC>2; p-value<0.05) is indicated.

**Supplementary Figure 6.**
